# Marine proteorhodopsins rival photosynthesis in solar energy capture

**DOI:** 10.1101/231167

**Authors:** Laura Gómez-Consarnau, Naomi M. Levine, Lynda S. Cutter, Deli Wang, Brian Seegers, Javier Arístegui, Jed A. Fuhrman, Josep M. Gasol, Sergio A. Sañudo-Wilhelmy

## Abstract

All known phototrophic metabolisms on Earth are based on one of three energy-converting pigments: chlorophyll-*a*, bacteriochlorophyll-*a*, and retinal, which is the chromophore in rhodopsins [1]. While the contribution of chlorophylls to global energy flows and marine carbon cycling has been studied for decades, the role of retinal-based phototrophy remains largely unexplored [1,2]. We report the first vertical distributions of the three energy-converting pigments measured along a contrasting nutrient gradient through the Mediterranean Sea and the Eastern Atlantic Ocean. The highest proteorhodopsin concentrations were observed above the deep chlorophyll-*a* maxima, and their geographical distribution tended to be inversely related to that of chlorophyll-*a*. We further show that proteorhodopsins potentially absorb as much or more light energy than chlorophyll-*a* –based phototrophy and this energy is sufficient to sustain bacterial basal metabolism. Our results suggest that ubiquitous proteorhodopsin-containing heterotrophs are important contributors to the light energy captured in the sea.

---

Sunlight drives virtually all life on the Earth’s surface, with about 50% of primary productivity occurring in marine systems [3]. However, prior to year 2000, all phototrophic metabolisms in the ocean were believed to be based on chlorophyll-like molecules. This traditional view of phototrophy changed radically with the discovery of marine bacterial rhodopsin-like proteins (proteorhodopsins: PRs) [4]. Metagenomic surveys indicated that PR occurs in ~75% of marine bacteria and archaea in oceanic surface waters [4,6,7]. However, it has been recently reported that >79% of marine bacteria from the Mediterranean Sea contain PRs while only <6% contain other types of rhodopsins (e.g., xantorhodopsins and actinorhodopsins) [8]. Metagenomic data from different types of environments have shown that PR gene abundance in <0.8 *µ*m microbial fractions exceeds the combined oxygenic and anoxygenic photosystem genes by threefold [8,9]. All of the available evidence strongly suggests that PR phototrophy is a vital and widespread process in all sunlit ecosystems [9,10]. However, PR quantification that relies on genes or transcripts does not necessarily reflect PR abundance, due to post-transcriptional regulation and other factors that result in a disconnect between gene transcription and protein abundance. Since the discovery of PRs, only a handful of estimates of PR abundance have been reported [11–13].

To try to establish the ecological relevance of PRs, we have modified the methods originally developed for haloarchaea [14,15] to quantify the light-sensitive pigment in rhodopsins, the chromophore retinal. X-ray crystallography data show that all rhodopsins (both microbial type I and metazoan type II) have a common structure: a typical seven transmembrane alpha-helical protein motif (opsin) binding a single molecule of retinal via a covalent Schiff base [16,17,18]. Therefore, because PRs have a single retinal chromophore associated with the polypeptide opsin, the total number of retinal molecules is equivalent to the total number of PRs [16,19]. Unlike proteomic approaches, the direct quantification of retinal assesses even previously uncharacterized PRs. Using this direct method, we have estimated the PR concentrations in microbial communities collected along an East-West transect in the Mediterranean Sea and the Eastern Atlantic Ocean (Figure 1A, Table S1). We also compared the abundance of PRs to the other two major pigments, chlorophyll-*a* (Chl-*a*) for oxygenic photosynthesis and bacteriochlorophyll-*a* (Bchl-*a*) for aerobic anoxygenic phototrophy (AAP), allowing us to estimate for the first time the distribution and contribution of each of these energy-transducing mechanisms to the total solar radiation captured in the surface ocean.

**Figure 1.**
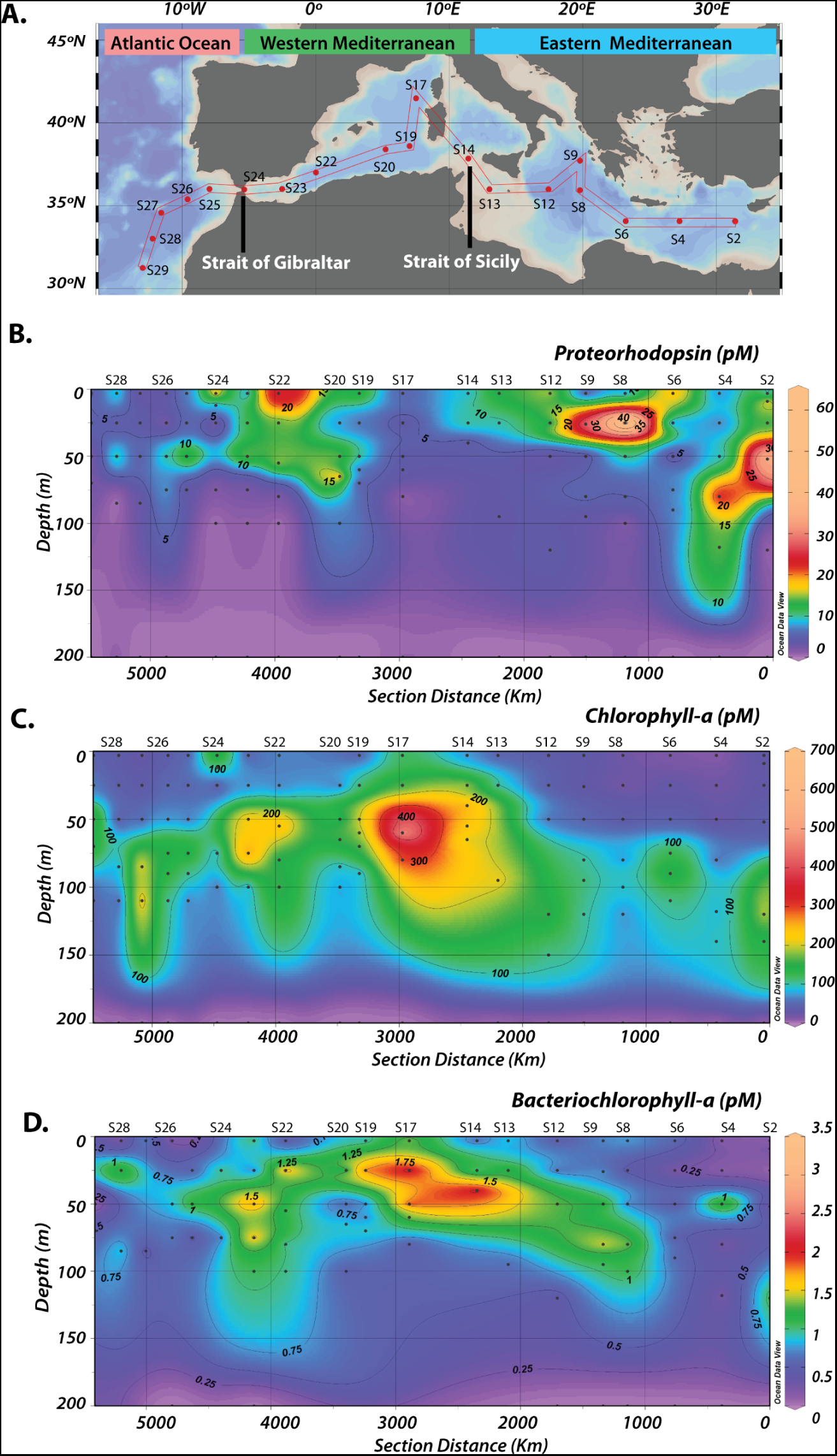
Sectional distributions of pigment concentrations measured along the Mediterranean Sea and in the Eastern Atlantic Ocean. **A)** Map of sampling stations, **B)** distribution of retinal in proteorhodopsins **C)** chlorophyll-*a*, and **D)** bacteriochlorophyll-*a*.The black symbols indicate the depths of sampling.

To better understand the processes influencing the different phototrophic mechanisms, we collected microbial planktonic samples at different locations representing very distinct oceanographic regimes: i) Oligotrophic areas in the Eastern Mediterranean sea (Stations 2-12), ii) Coastal influenced regions in the Western Mediterranean sea, (Stations 13-24), and iii) Open ocean environments in the Eastern Atlantic ocean (Stations 25-29; Figure S1AB). The highest PR concentrations were observed above the deep chlorophyll maximum in almost all stations (Figure 1BC, Figure 2, Table S1). This may relate to the light wavelengths maximally absorbed by PRs, which attenuate more rapidly in the water column than those maximally absorbed by Chl-*a* [11,20]. The highest PR concentrations were observed in the low nutrient regions of the Eastern Mediterranean Sea, with maximum values ranging from 14.4 to 61.2 pM (stations 2-12). This trend contrasts with the geographical distribution of Chl-*a*, which peaked in the most coastal influenced regions (68.7-695.6 pM; stations 13-24). Therefore, the longitudinal distribution of PR photoheterotrophy showed highest values in areas where oxygenic photosynthesis was the lowest. Bchl-*a* concentrations were about an order of magnitude lower than those of PR throughout the entire transect (0.5-3.5 pM), reflecting the overall low abundance of photoheterotrophic AAP bacteria in the Mediterranean Sea [21,22]. As previously observed [22], the highest levels of Bchl-*a* were measured in the coastal-influenced regions of the Western Mediterranean (Figure 1D, Figure 2). Thus, our data suggests that PR is the dominant photoheterotrophic pigment, on a molar basis, exceeding Bchl-*a* by an order of magnitude. Moreover, the observed pigment distributions support the hypothesis that PR phototrophy is particularly relevant in oligotrophic environments, where organic nutrients are scarce [23–27]. Notably, we did not detect PR or Bchl-*a* in the >3 μm pore-size fraction of any of our samples, even though rhodopsin genes and transcripts have previously been found in large phytoplankton [28,29]. This suggests that PR-dependent metabolisms were less important for protists (or for particle-attached prokaryotes) than they were for the picoplanktonic communities in these regions. The lack of PRs in the large pore size fraction during our study also suggests that viral-like rhodopsins were not present or they were in very low abundances, as the only viral proteorhodopsin genes identified to date belong to viruses that infect protists [30].

**Figure 2.**
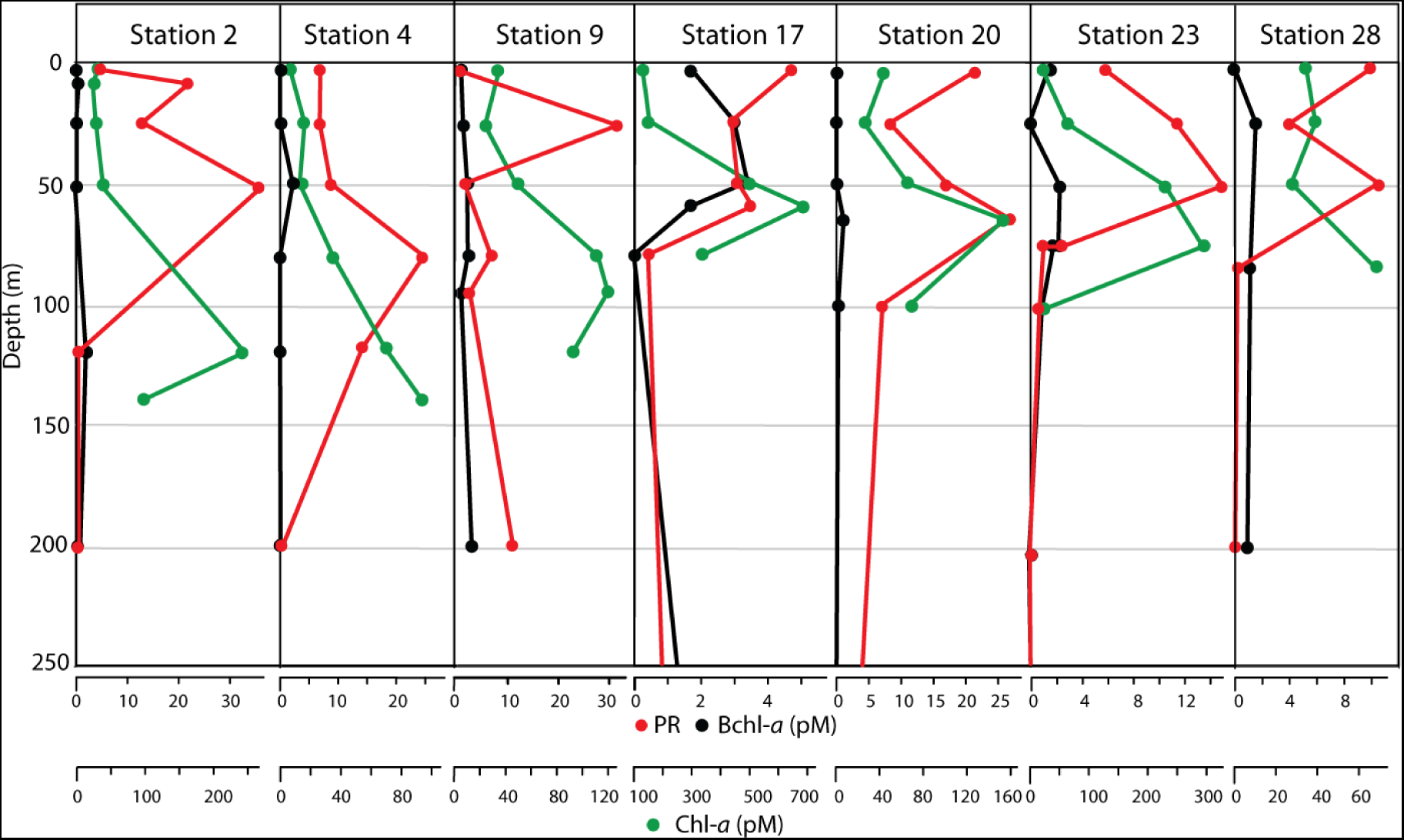
Depth-profiles of pigment concentrations (PR, proteorhodopsin; Bchl-*a*, bacteriochlorophyll-*a*; Chl-*a*, chlorophyll-*a*) measured at the different basins of the Mediterranean Sea and Eastern Atlantic Ocean. Stations 2, 4 and 9 are representative of oligotrophic regions of the Eastern Mediterranean; stations 17, 20 and 23 were sampled in the coastal-influenced Western Mediterranean and station 28 in the Eastern Atlantic Ocean.

Although the PR concentrations measured along the oceanographic transect were about an order of magnitude lower than the levels of Chl-*a* (Figure 1), the light-harvesting and energy transfer capabilities of each photosystem is not a function of the absolute pigment concentrations but depends on the number of pigment molecules cluster to form each “photosynthetic” unit (PSU) (300 molecules for Chl-*a*; 34 for Bchl-*a*; 1 for PR) [18,31,32]. Along the Mediterranean-Atlantic cruise, PRs have the highest number of PSUs (1×10^11^-4×10^13^ PSUs L^−1^), exceeding the amount of PSUs of Chl-*a* by an order of magnitude (1×10^10^-1×10^12^ PSUs L^−1^) (Table S1). The number of PSUs for Bchl-*a* was several orders of magnitude lower than the PSUs calculated for the PR and Chl-*a* (2×10^9^-7×10^10^ PSU L^−1^; Table S1). The amount of depth-integrated energy potentially captured by retinal-based PR, Chl-*a*-based photosynthesis, and Bchl-*a*-based AAP in the water column (in J m^−2^ sec^−1^; Figure 4, Table S1) reflect the trends observed in the number of the different photosynthetic units measured along the cruise. We found that PR could capture the highest amount of radiant energy throughout the entire transect (ranging from 0.32 to 3.09 J m^−2^ sec^−1^). Those estimates are about an order of magnitude higher than the cumulative or depth-integrated solar energy captured by oxygenic photosynthesis by autotrophic phytoplankton in the Eastern Mediterranean (Chl-*a*-based energy range: 0.05-0.16 J m^−2^ sec^−1^). In contrast, the extent of solar radiation absorbed by AAP bacteria (0.001-0.021 J m^−2^ sec^−1^) was orders of magnitude lower than that absorbed by the organisms containing the other two photosystems. These patterns were consistent regardless of whether the pigment specific absorbance wavelengths or the total photosynthetically available radiant energy (PAR) were used in our calculations (Figure 2).

The spatial gradient of cellular PR quotas (PR molecules cell^−1^) measured along our transect suggests that PR photoheterotrophy is particularly favored in the nutrient-limited areas of the Mediterranean (Figure 3). For example, the cellular PR content measured in the oligotrophic picoplankton from the Eastern Mediterranean ranged from 38,000 to 140,000 molecules cell^−1^, exceeding by almost an order of magnitude earlier estimates [2]. Similar rhodopsin quotas have only been observed in Haloarchaea species from hypersaline environments [33]. In contrast, cellular PR content from the Eastern Atlantic Ocean (5,000-14,000 PR molecules cell^−1^) and the Western Mediterranean Sea (4,000-42,000 PR molecules cell^−1^) were consistent with previous measurements in both environmental and pure culture samples. Thus, the spatial distribution in the PR quotas seems to relate to inorganic nutrient concentrations (Figure S1), suggesting that the characteristics of different nutrient regimes might have a profound impact on the type of metabolism (heterotrophy *vs*. phototrophy) used by photoheterotrophic bacteria.

**Figure 3.**
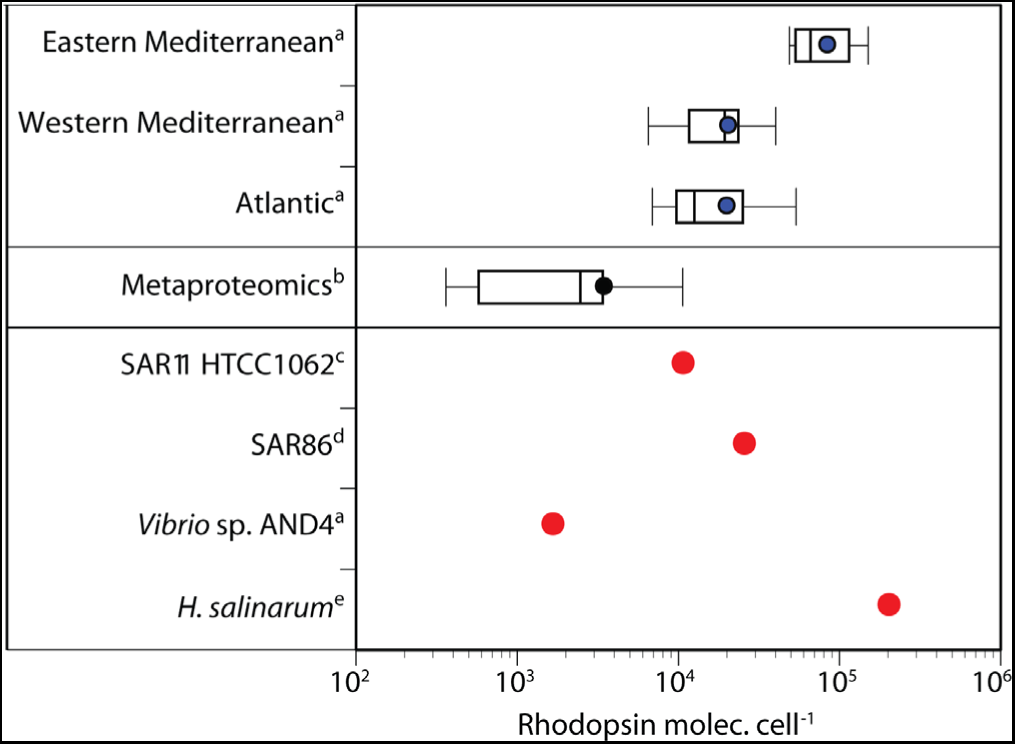
Rhodopsin cellular quotas (molecules cell^−1^) measured in the picoplankton collected in the different basins of the Mediterranean Sea and in the Eastern Atlantic Ocean compared to prior studies. ^a^Samples measured in this study: Eastern Mediterranean (stations 2-12),Western Mediterranean (stations 13-24), and Atlantic Ocean (stations 25-29) and Vibrio sp.AND4; ^b^metaproteomics estimates from [2]; laser flash photolysis measurements: ^c^SAR11 bacterium Candidatus Pelagibacter ubique HTCC1062 [12], ^d^SAR86 bacteria [11], ^e^Halobacterium salinarum [33].The line within the box-plot is the median, the dot is the mean and the boundary of the boxes indicates the 25th and 75th percentiles. Error bars to the left and to the right of the box indicate the 10th and 90th percentiles.

To evaluate the possible physiological advantages provided by photoheterotrophy to PR-containing bacteria, we estimated the potential highest energy yields on a per cell basis provided by this photosystem (in kJ cell^−1^ day^−1^; Figure 5). The highest cellular energy yield from PR ranged from 1 × 10-13 to 2 × 10-12 kJ cell^−1^ day^−1^ along the Mediterranean Sea-Atlantic Ocean transect. Those PR-energy per cell yields were between 1 to 4 times lower and between 2 to 57 times lower than the energy provided by Chl-*a* (2-18 × 10^−12^ kJ cell^−1^ day^−1^) in the Eastern and Western Mediterranean respectively. However, with far more PR-containing cells than chlorophyll-containing ones, the cumulative or depth-integrated energy capture from PR was larger, as shown above (Figure 4). Overall, the observed geographical trends support the notion that PR photoheterotrophy is a particularly vital mechanism for bacterioplankton survival in areas where other resources are scarce [23–27]. So while PR-containing taxa are found in low and high nutrient concentration waters worldwide [5], this mechanism is exceptionally valuable in sunlit oligotrophic locations, possibly including the South Pacific Gyre, an area much larger than the Eastern Mediterranean.

**Figure 4.**
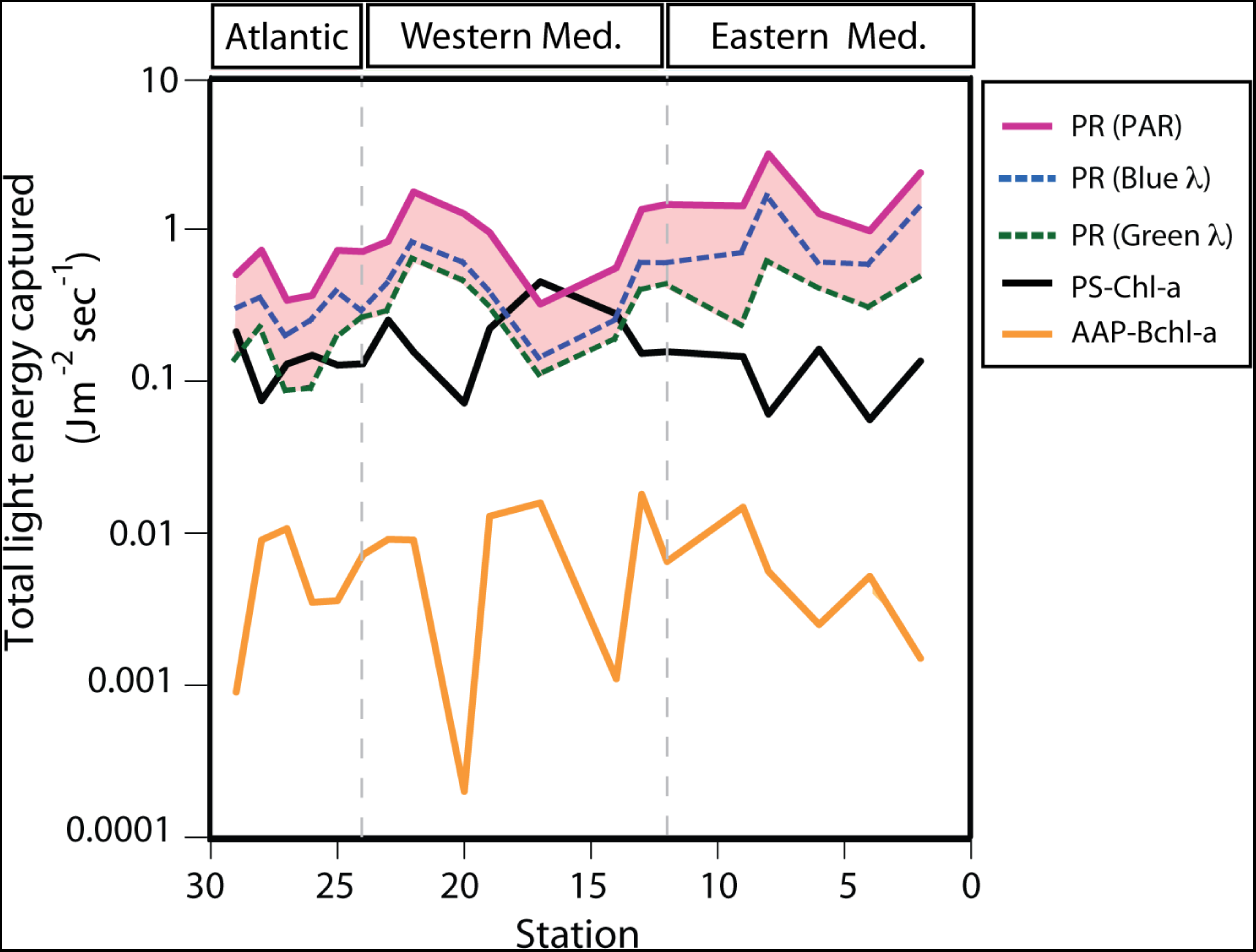
Geographical distribution of the total light energy potentially captured by proteorhodopsin (PR), Chlorophyll-*a* photosystem (PS-Chl-*a*), and aerobic an-oxygenic phototrophy (AAP-Bchl-*a*) in the Mediterranean Sea and Eastern Atlantic Ocean. These data represent the daily average depth-integrated energy absorbed in the water column.A photocycle of 10 ms was used for the PR calculations, as DNA sequence, spectral tuning and kinetics data[6,39] show that most PRs from surface temperate waters have fast photocycles typical of H+ pumps. Solid lines denote estimates using all photosynthetic available radiation (PAR; 400-700 nm). They represent the maximum energy estimates, which assume that also accessory pigments can access all wavelengths (PAR; 400-700 nm). Conservative energy calculations using specific wavelengths for blue-absorbing (490 nm) and green-absorbing (530 nm) PR are shown in dashed lines.

**Figure 5.**
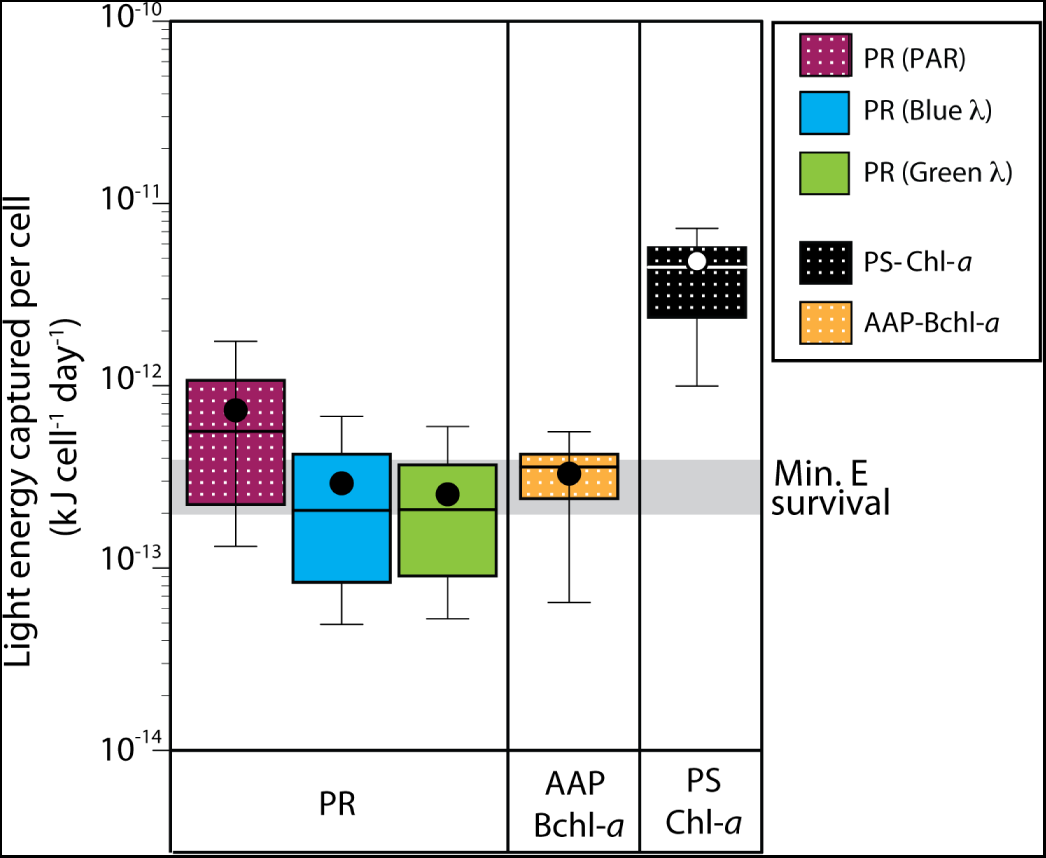
Light energy captured per cell by proteorhodopsin (PR), bacteriochlorophyll-*a* (AAP-Bchl-*a*) and chlorophyll-*a* (PS-Chl-*a*) in the Mediterranean Sea and Eastern Atlantic Ocean. These calculations assumed hyperbolic response to light, that 75% of the total heterotrophic bacteria contain PR genes and 2.5% contain Bchl-*a* (Supplementary Material and Methods and Table S6). Chl-*a* containing cells were estimated as the addition of the Prochlorococcus, Synechococcus and picoeukaryote counts measured with flow cytometry.As reference, the grey horizontal band denotes the estimated energy range necessary to maintain basal metabolism or survival in heterotrophic bacteria [2]. Dotted boxes denote estimates using all photosynthetic available radiation (PAR; 400-700 nm). Conservative calculations using only specific wavelengths for blue-absorbing (490 nm) and green-absorbing (530 nm) PR are shown as blue and green boxes.

Our calculations also showed that the cellular energy yield captured by PR was within the same order of magnitude of the minimum survival energy threshold for some heterotrophic bacteria (2 to 4×10^−13^ kJ cell^−1^ day^−1^) [2]. However, at some locations in the Eastern Mediterranean, the energy acquired by PR was above that threshold by an order of magnitude (Figure 5). Thus, our results suggest that PR alone provides, at least, enough energy to each individual bacterium, on average, to sustain its basal metabolism. From the perspective of photoheterotrophy, our per-cell energy calculations are consistent with the fact that known AAP- and PR-containing bacteria are still heterotrophs and require organic matter for biomass synthesis or growth. These data also support the hypothesis that the most extremely oligotrophic marine environments (e.g. Eastern Mediterranean) have such low primary production from photosynthesis that complementary light-dependent metabolisms can supply as much or possibly more energy to heterotrophic prokaryotes than they can obtain from organic matter through classic food web based energy transfers.

Finally, although we were not able to determine how much of the harvested solar energy is transformed into actual biological functions such as light-enhanced substrate uptake [27] or ATP synthesis [26], our results suggest that PR-photoheterotrophy captures and potentially channels significant amounts of radiant energy to the biosphere. Future challenges will include understanding to what extent and in which manner this prevailing type of phototrophy affects the processing of organic matter and therefore its role in the marine carbon cycle. The observed large contribution of PRs to the energy budget in less productive regions suggests that retinal-based phototrophy is likely to increase as oligotrophic areas continue to expand in the world ocean in response to climate change forcing [34].

## Methods

Seawater samples were collected along a Mediterranean Sea-Atlantic Ocean transect (Figure 1) on board the Spanish ship RV Sarmiento de Gamboa, from April 30^th^ to May 28^th^, 2014. Sampling dates and station coordinates are given in Table S1. Samples were collected at various depths within the photic zone using a CTD rosette equipped with 12 – liter Niskin bottles. For the retinal and Bchl-*a* analyses, 8 liters of seawater were filtered per depth using a peristaltic pump connected to a sequential filtration system. To avoid the presence of large planktonic organisms in the samples, water was pre-screened using a nylon 10-μm mesh attached to the inflow of the filtration system. The 3-10μm and the 3μm-0.2μm microbial size fractions were collected using 47 mm, 3μm pore-size polycarbonate filters (Millipore corp., Billerica, MA) and Sterivex-GV filter units (Millipore), respectively. Filters were stored at −80ºC until extraction. The retinal extraction and analysis protocol was modified from the literature [14,15], and optimized for marine samples using cultures of the PR-containing bacterium *Vibrio* sp. AND4 and the knockout mutant strain AND4-Δprd [24] (Supplementary Information). Briefly, the filters were sonicated in 3 ml of LC-MS grade methanol on ice and in the dark for 30 seconds. 250 μl of 1% butylated hydroxytoluene (BHT) were subsequently added, and the samples were left to extract for 24h at −20ºC. After the extraction, a 0.5 ml aliquot was used for Bchl-*a* analysis by HPLC [34]. 100µl of 1 M hydroxylamine was added to the remaining sample extraction and was incubated under light for 2 hours. PR concentration was determined as retinal oxime using liquid chromatography-mass spectrometry (LC-MS). The technique is fully described in the Supplementary Information. For Chl-*a* measurements, 500 ml of seawater were filtered through 25 mm GF/F filters (Whatman) using low vacuum. The filters were frozen at −20 ºC before pigments were extracted on board in 90% acetone for 24 h in the dark at 4 ºC. Chl-*a* concentrations were estimated by fluorometry in a Turner Designs fluorometer calibrated with pure Chl-*a* (Sigma Chemical)[36]. Dissolved inorganic nutrients (nitrate, phosphate) were collected from the Niskin bottles in 20-ml acid-washed polyethylene flasks and determined on board using standard segmented flow analysis methods with colorimetric detection measured on board [37]. Autotrophic picoplankton (Prochlorococcus, Synechococcus, and picoeukaryotes) and heterotrophic prokaryotes were enumerated by flow-cytometry (Becton– Dickinson FACScalibur) [38]. The bio-energetic calculations used to determine the radiant energy captured by each photosystem are described in the Supplementary Material and Methods.

## Acknowledgements

We thank X.A.Alvarez Salgado and V.Vieitez for the inorganic nutrient data, the crew on board the vessel Sarmiento de Gamboa for sample collection, Erin Fichot and Christopher Suffridge for providing seawater samples for method optimization.This project was founded by the Marie Curie Actions– International Outgoing Fellowships (project 253970), the US National Science Foundation grant OCE1335269 and the Gordon and Betty Moore Foundation Marine Microbiology Initiative award 3779. J.A. and J.M.G were supported by the Spanish project HOTMIX (CTM2011-30010-C02).We thank Rodolfo Iturriaga for comments and relevant discussions.

## Author contributions

LG-C and SS-W designed the study. LG-C, SS-W and JMG planned the sampling locations and ancillary parameters to be measured during the cruise. SS-W, LG-C, LSC and BS contributed to the development of the retinal oxime analytical technique and DW to the bacteriochlorophyll-*a* measurements. NML and DW analyzed and did calculations for the solar energy capture data. JA was the scientific chief during the Mediterranean-East Atlantic Ocean cruise and provided with Chlorophyll-*a* and flow cytometry data. LG-C, NML, LSC, DW, BS, JA, JAF, JMG and SS-W analyzed data and contributed to the writing of the manuscript.

